# 3D Nuclei Segmentation by Combining GAN Based Image Synthesis and Existing 3D Manual Annotations

**DOI:** 10.1101/2023.12.06.570366

**Authors:** Xareni Galindo, Thierno Barry, Pauline Guyot, Charlotte Rivière, Rémi Galland, Florian Levet

**Author notes:** These two authors contributed equally (Xareni Galindo, Thierno Barry).

## Abstract

Nuclei segmentation is an important task in cell biology analysis that requires accurate and reliable methods, especially within complex low signal to noise ratio images with crowded cells populations. In this context, deep learning-based methods such as Stardist have emerged as the best performing solutions for segmenting nucleus. Unfortunately, the performances of such methods rely on the availability of vast libraries of ground truth hand-annotated data-sets, which become especially tedious to create for 3D cell cultures in which nuclei tend to overlap. In this work, we present a workflow to segment nuclei in 3D in such conditions when no specific ground truth exists. It combines the use of a robust 2D segmentation method, Stardist 2D, which have been trained on thousands of already available ground truth datasets, with the generation of pair of 3D masks and synthetic fluorescence volumes through a conditional GAN. It allows to train a Stardist 3D model with 3D ground truth masks and synthetic volumes that mimic our fluorescence ones. This strategy allows to segment 3D data that have no available ground truth, alleviating the need to perform manual annotations, and improving the results obtained by training Stardist with the original ground truth data.

## 1 INTRODUCTION

The popularity of 3D cell cultures, such as organoids or spheroids, has recently exploded due to their ability to offer valuable models to study human biology, far more physiologically relevant than 2D cultures (Jensen and Teng, 2020; Kapałczyńska et al., 2018). Nowadays, automatically acquiring hundreds of organoids in 3D has become a reality thanks to the advances in microscopy systems (Beghin et al., 2022). Life scientists have therefore access to distributions of nuclei in 3D for a wide diversity of cell types and growing conditions, a key feature forming the basis of advanced quantitative analysis of important cell functions. However, 3D cellular cultures inherently display a large diversity of nuclei features, shapes and arrangements. And 3D microscopy of such complex samples most often leads to images with lower contrast and/or signal to noise ratio as compared to 2D microscopy of single cellular layer, with overlapping nuclei according to the achieved optical sectioning. Hence, accurate and automated nuclei segmentation in these conditions has turned out to be highly complex. The bottleneck has therefore shifted from the acquisition to the downstream analysis and quantification steps.

Many computational solutions have been proposed over the years that uses traditional image processing methods to tackle this segmentation problem (Caicedo et al., 2019; Malpica et al., 1997; Li et al., 2007), in 2D and 3D. However, they are usually tailored for a specific application and do not generalize well, resulting in the necessity to adapt their parameters and ultimately preventing an automatic and bias-free analysis. In parallel, the last few years have witnessed the rapid emergence of convolutional neural networks (CNNs) as a method of choice for microscopy image segmentation, achieving an accuracy unheard of for a wide range of segmentation tasks. Among those, deep learning-based nuclei segmentation has been particularly active, with more than one hundred methods being developed since 2019 (Mougeot et al., 2022).

Supervised approaches, in which paired of images and masks are used for training, are the current state of the art for nuclei segmentation, with Stardist (Schmidt et al., 2018; Weigert et al., 2020) and Cellpose (Stringer et al., 2021) having reached a prominent position. Their adoption has been facilitated by their integration into several open-source platforms (von Chamier et al., 2021; Gómez-de Mariscal et al., 2021; Sofroniew et al., 2022), in which life scientists are able to directly use several pre-trained models (Caicedo et al., 2019) to segment their images. Unfortunately, these pre-trained models are limited to the segmentation of nuclei in 2D, a fact directly related to the lack of available ground truth (GT) datasets in 3D. Determining the organization of nuclei in 3D therefore requires life scientists to annotate themselves their data, a daunting and time-consuming task which turns out to be much more challenging than in 2D due to the lower image quality and more crowded cell populations.

Generating synthetic training datasets has therefore emerged as a potential solution to this problem through the use of conditional Generative Adversarial Networks (cGANs) (Goodfellow et al., 2014; Isola et al., 2018; Zhu et al., 2017). cGANs learn a mapping from an observed source image *x* and a random noise vector *z*, to a “transformed” target image *y, G* : *x, z* → *y* (Fig. 1). For nuclei segmentation, the use of such generative networks aims to generate realistic microscopy images (background, signal to noise ratio, inhomogeneity, …) of nuclei (target style) from binary masks (source style) by training the cGAN with paired or unpaired datasets (Baniukiewicz et al., 2019; Fu et al., 2018; Wang et al., 2022). Nevertheless, GANs are notoriously known to be difficult to train, with a large number of hyperparameters required to be tuned.

**Figure 1:**
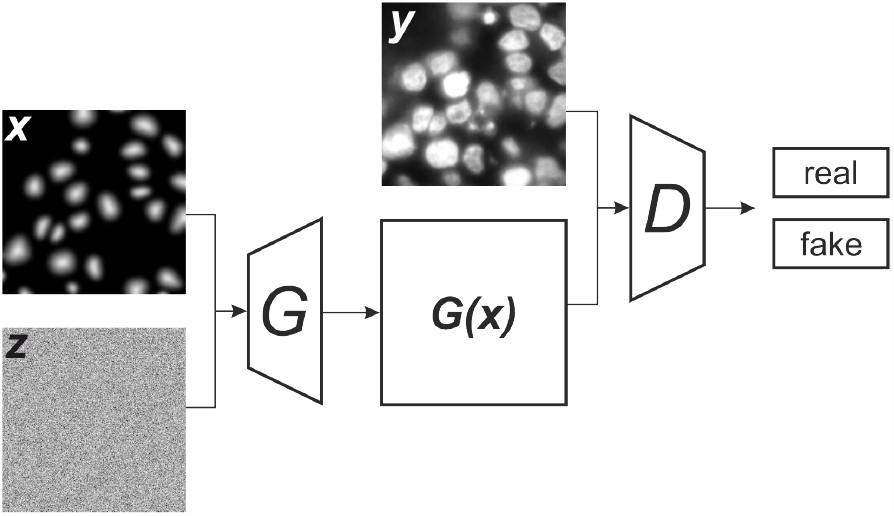
Architecture of FCGAN. In addition to an image in the source style, the generator takes a noise image as input. The discriminator meant to determine if the generated synthetic image *G*(*x*) is real or fake is multi-scale. Please note that the FCGAN was trained on the whole images, and not on crops as shown in this illustration.

In this work, we present a workflow to segment nuclei in 3D when no specific GT exists by leveraging on cGANs to generate fluorescence synthetic volumes of nuclei from 3D masks. While life scientists start to be accustomed to using packaged deep learning methods for segmentation or classification, image synthesis is still far from being accessible to regular users. We therefore made the choice to specifically design our workflow to be accessible to nonexpert life-scientists, leveraging on already packaged methods, both for the segmentation and the image generation, with the aim to avoid any tedious hand-annotating steps. With that in mind, our workflow first relies on the segmentation of the individualized 2D planes of our acquired volumes using the already pre-trained, well established and now robust 2D segmentation model Stardist 2D *db*2018 (Schmidt et al., 2018). After modification of the instance masks by applying a distance transform and a Gaussian filter, we pair these transformed ‘GT’ masks with the fluorescent nuclei images to train a cGAN specifically designed for biological images, that do not require complex hyper-parameters tuning or specific cropping data selection (Han et al., 2020). It allows to generate 3D volumes of nuclei, that resemble the fluorescent volume we acquired, from existing 3D GT masks that were typically created for a different cell type or microscopy modality. Finally, those pairs of GT mask volumes and synthetic fluorescent volumes of nuclei are used to train a Stardist model in 3D dedicated to our 3D samples type and imaging modality.

Through this workflow, we demonstrate that using existing works to their full extent can (i) circumvent the requirement of tedious hand-annotating steps for the segmentation of nuclei in 3D in complex and crowded environment, and (ii) alleviate the need to develop new deep learning architectures in an already crowded field (Mougeot et al., 2022), while also facilitating their usage by life scientists. We applied this workflow on a couple of cell lines and microscopy modalities and show that we managed to have good qualitative segmentation results without having to specifically hand annotate new volumes. This approach greatly widens the scope of use of 3D segmentation methods in the rapidly growing and diversifying field of 3D cell culture analysis, where many imaging modalities are explored and cell types with their own morphology characteristics used.

## 2 MOTIVATION

We recently acquired several oncospheres in 3D of the colorectal cancer cell line HCT-116 (Goodarzi et al., 2021) expressing the nucleus fluorescent label Fucci with a confocal microscope, denoted as *I*^*c*^, that we aim to segment while not having any GT mask for that cell culture type and imaging modality. In parallel, we were kindly provided a pre-trained Stardist 3D model 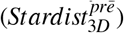 with the corresponding GT dataset by the authors of (Beghin et al., 2022). The GT dataset was composed of 7 volumes that were manually annotated and that we will denote as 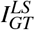 and 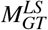 for the images and masks, respectively. These 7 volumes were composed of different 3D cell culture types (oncospheres and neuroectoderms), nuclear staining (DAPI and SOX) and z-steps (500 nm and 1 μm). In addition, they have been acquired with the soSPIM imaging technology (Galland et al., 2015), a single-objective light-sheet microscope different from the confocal imaging modality used to acquire the oncospheres we aimed to segment. With this 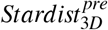 model, the auhtors of (Beghin et al., 2022) managed to automatically segment more than one hundred oncospheres and neuroectoderms acquired with the soSPIM imaging modality. However, applying 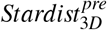 to *I*^*c*^ gave poor results, as it only managed to identify a portion of the nuclei with an overall oversegmentation, most probably due to the different cell morphology and imaging modality of *I*^*c*^ as compared to the GT datasets used to train the model.

We therefore tried to generate synthetic volume having the same style than *I*^*c*^ with SpCycleGAN (Fu et al., 2018), with the final objective of being able to train a new Stardist 3D model more adapted to our data. SpCycleGAN allows to generate synthetic volumes from binary masks without GT by being built on top of the unpaired image-to-image translation model CycleGAN (Zhu et al., 2017). Being unpaired, the network can use *I*^*c*^ as target style without requiring the corresponding nuclei segmentation as source. Originally, the authors of (Fu et al., 2018; Wu et al., 2023) generated 3D nuclei masks by filling volumes with binarized (deformed) ellipsoids even though it could be limiting as it may not faithfully represent the nuclei morphology. Having already a GT nuclei masks, we therefore decided to train SpCycleGAN with 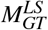and *I*^*c*^ to avoid this pitfall.

In our hand, SpCycleGAN failed to generate realistic microscopy images (Fig. 2). Scrutinizing the toy datasets available with the network, we tested different scaling and cropping of the data, as well as removing any crops having void regions during training. Unfortunately, none of our tests were conclusive. We hypothesize that this may be related to two reasons. First, volumes composing *I*^*c*^ exhibit a high level of noise and an overall crowded and overlapping nuclei population. SpCycleGAN may fail to transfer all these features from binary masks. Second, SpCycleGAN may require parameters fine-tuning to work properly, but this goes against our objective of having a more generic workflow that could be used by nonexperts.

**Figure 2:**
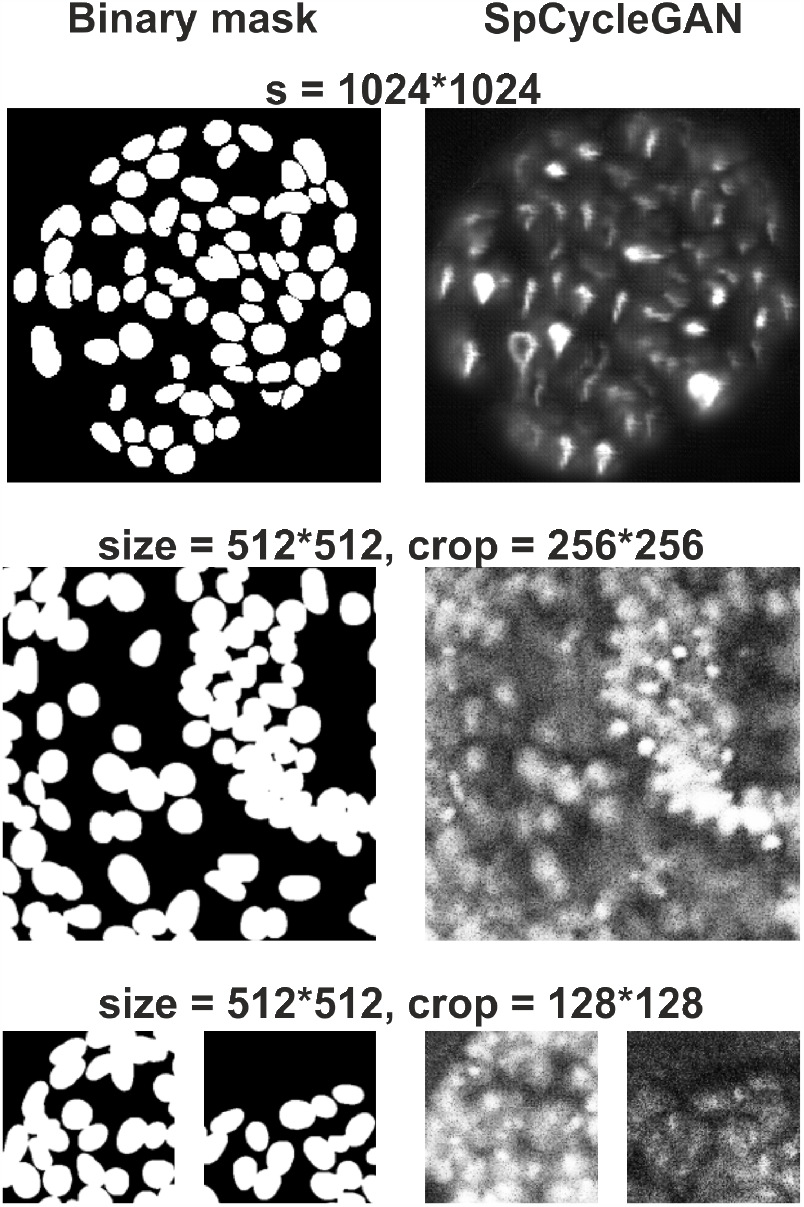
SpCycleGAN fails to generate realistic microscopy images even with different scaling and cropping.

## 3 METHOD

The proposed method is based upon the use of well established and more robust deep-learning methods for nuclei segmentation (Stardist (Schmidt et al., 2018; Weigert et al., 2020)) and image synthesis (conditional GANs in a slightly modified version (Han et al., 2020)). Ours choices are motivated by the fact that we wanted an as-simple-as-possible backbone. In this manner, we intend to show that a simple approach focusing on data manipulation combined with existing networks can perform qualitatively well and propose a workflow that could become generic to enable the segmentation of a large variety of 3D biological samples acquired upon different imaging modalities.

### 3.1 Fully-Conditional GAN

Contrary to natural images in which every local region contains relevant information, biological images are composed of a mix of informative and void regions. They are also inherently multiscale with both the large-scale spatial organization of cells and their individual morphology and texture being essential. Since GANs have been originally developed for natural images, they tend to struggle to capture these intertwined features and to fail to generate realistic void regions.

We therefore based our synthesis method in the fully-conditional GAN (FCGAN) architecture (Han et al., 2020), an improvement of GAN focused on modifications that allow to synthesize multi-scale biological images (Fig. 1). This architecture consists of a generator built upon the cascaded refinement network (Chen and Koltun, 2017) instead of the more traditional U-Net (Ronneberger et al., 2015; Isola et al., 2018) architecture as it is less prone to mode collapse. They also added two modifications to the traditional GAN architecture. First, they modified the input noise vector *z* to a noise “image”, i.e. a 3D tensor with the first two dimensions corresponding to the spatial positions. Instead of a noise vector that limits the size of output images, modifying the noise image size allows to output synthetized images of arbitrary sizes. Second, they used a multi-scale discriminator. Having a fixed size discriminator limits the quality and coherency of synthetized images to a micro (object) or macro (global organization) level. Using a multi-scale discriminator ensure the generator to produce images both globally and locally accurate, a desired feature when dealing with biological images that exhibit several levels of organization. All these modifications allow to directly train the FCGAN with the acquired images, alleviating the need to fine-tune the parameters and resize or crop the void regions.

### 3.2 Image Synthesis Based 3D Stardist Training

In spite of its numerous advantages, FCGAN requires however paired images for training, therefore compelling us to provide corresponding mask and image pairs. Fortunately, 2D pre-trained models of well-established nuclei segmentation methods such as Stardist (Schmidt et al., 2018) or Cellpose (Stringer et al., 2021) have been trained with thousands of 2D GT masks and achieve now a high degree of robustness over a large variety of sample types. Consequently, we can directly determine the nuclei spatial distribution by segmenting each of the 2D frame (1024 1024) of the volumes of *I*^*c*^ with the 2D pretrained *db*2018 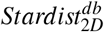 model (Fig. 3) instead of simulating them. These 2D segmentation will be denoted 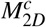 .

**Figure 3:**
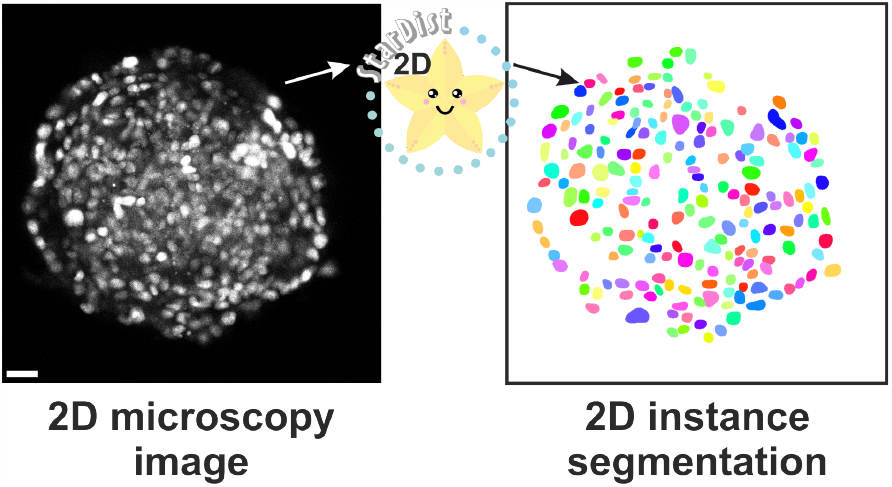
Nuclei segmentation with the 2D *db*2018 pretrained Stardist model results in sufficiently good results to train a conditional GAN, scale bar = 20 μm.

For image generation through the FCGAN model, binary and instance masks can lead to fuzzy results because their gradient can be too rough during backpropagation (Long et al., 2021) (Fig. 4(a)). We therefore applied a distance transform to our instance masks (Long et al., 2021) followed by an intensity normalization and a Gaussian filter (see Source code 1), which allows to remove intensity stiff jumps and dependency to the nuclei length, ultimately facilitating the style transfer performed by the FCGAN model (Fig. 4(b)).

**Figure 4:**
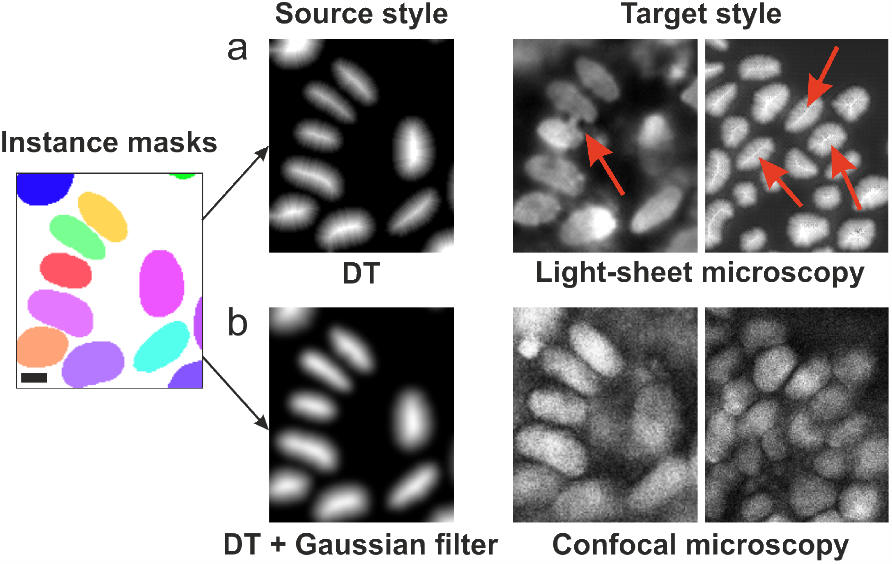
Manipulating instance masks to generate the FCGAN’s source style, scale bar = 5 μm. (a) After applying a distance transform on the binarized instance masks, the FCGAN generates plausible images but with visible artefacts related to the rough gradient of the transform (red arrows). (b) Applying a Gaussian filter after the mask binarization helps the FCGAN to learn a proper mapping between the source and target style.

**Source code 1:**
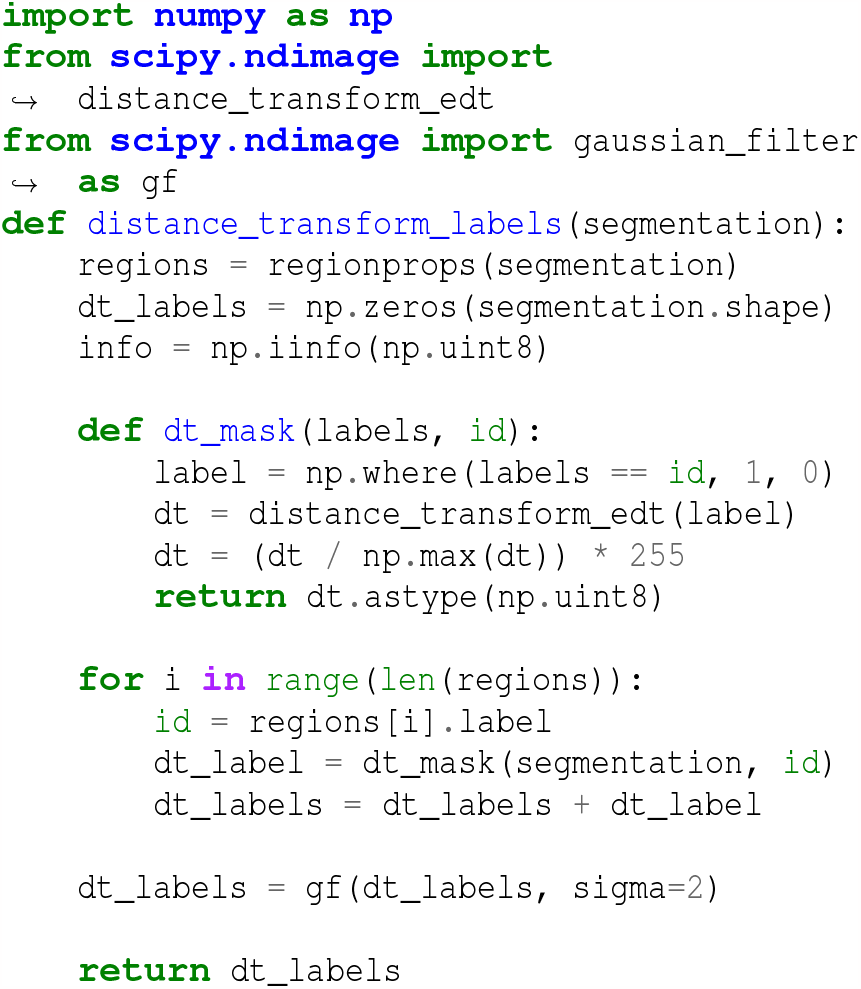
Transforming instance masks to the FCGAN’s source style.

The function described in Source code 1 was applied on each mask image of 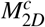, to obtain a set of transformed mask images 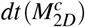 . Consequently, pairing 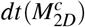 (source style) with the corresponding images from *I*^*c*^ (target style), allowed us to train the *FCGAN*^*c*^ model.

Finally, we generated 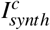, synthetic confocalstyled volumes (Fig 5) obtained by applying *FCGAN* to the transformed masks 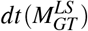 of the 3D GT dataset 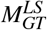. This was carried out after ensuring that the sizes of the masks composing 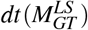 were similar to the sizes of the nuclei present in *I*^*c*^ . We then trained a 3D Stardist model called 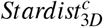 with the paired 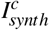 volumes and the corresponding 3D masks 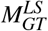 .

**Figure 5:**
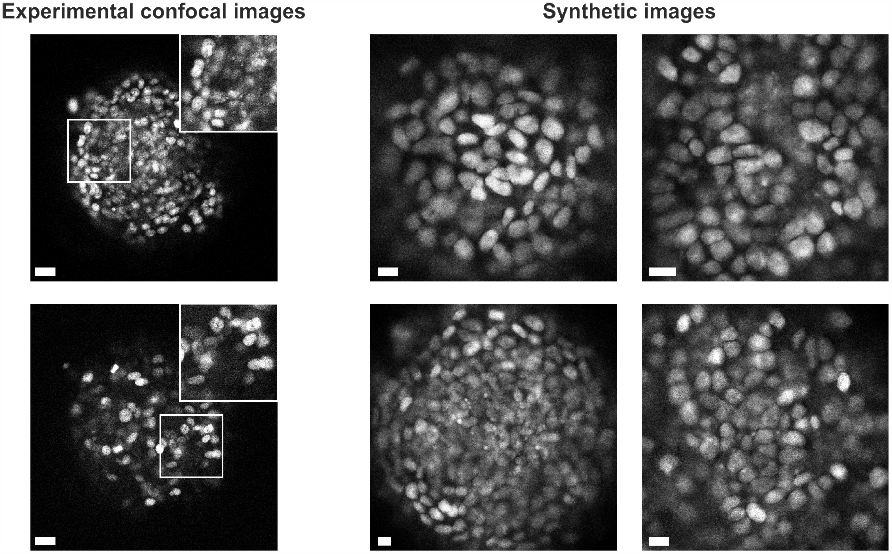
Comparison of experimental confocal images (scale bar = 20 μm) and synthetic images generated with the FCGAN style transfer (scale bar = 10 μm).

## 4 RESULTS

To assess both qualitatively and quantitatively the segmentation accuracy of our workflow, we compared it to two baselines and used two different cell cultures acquired with different microscopy modalities. In addition to *I*^*c*^, we thus acquired with a spinning-disc confocal microscope 4 suspended S180-E-cad–GFP cells (Chu et al., 2004) spheroids which nuclei were stained with DAPI (denoted as *I*^*SD*^). Indeed, the volumes composing *I*^*c*^ are highly challenging because of their crowded overlapping nuclei population and the overall level of noise. This makes them unsuitable to evaluate the segmentation accuracy of our workflow other than qualitatively, since it is even difficult by eye to determine the limits of each nucleus. On the contrary, *I*^*SD*^ were composed of a lower number of cells over a shorter thickness (5-10 μm) ensuring to have a sparser distribution of nuclei per volume (90 nuclei for the 4 volumes) easier to asses. On the side of the segmentation, and in addition to 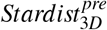, the second baseline was obtained by training a 3D Stardist model 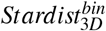 to learn how to reconstruct 3D instance masks from binarized masks, i.e. by pairing 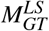 with its binarization. 3D segmentation with 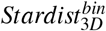 were therefore done by applying the model directly on the binarized 2D masks obtained with the pre-trained 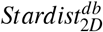 model on *I*^*c*^ and *I*^*SD*^.

### 4.1 Qualitative Assessment with *I*^*c*^

Applying 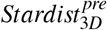 (Beghin et al., 2022) gave poor results with the volumes composing *I*^*c*^ for two reasons (Fig. 6). First, this model was trained with masks smaller than the nuclei present in *I*^*c*^, resulting in oversegmentation. Second, due to the different imaging modality, *I*^*c*^ exhibited more noise with its nuclei having a different texture than the ones from 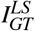 . It resulted with 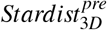 missing a lot of nuclei when applied to *I*^*c*^. On the other hand, 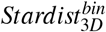 managed to segment more nuclei than 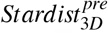 .since the 2D segmentation provided by 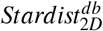 managed to identify more nuclei on a per frame basis. Nevertheless, results shows that 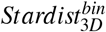 struggles to handle isolated 2D masks, resulting in over-segmented small nuclei (Fig. 7 and Fig. 6). On the contrary, the segmentation provided by 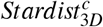 gave better qualitative results than 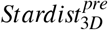 and 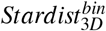 (Fig. 6). In particular, 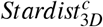 managed to segment nuclei exhibiting a wide range of intensity values, from dim to intense, while preventing over-segmentation at the same time.

**Figure 6:**
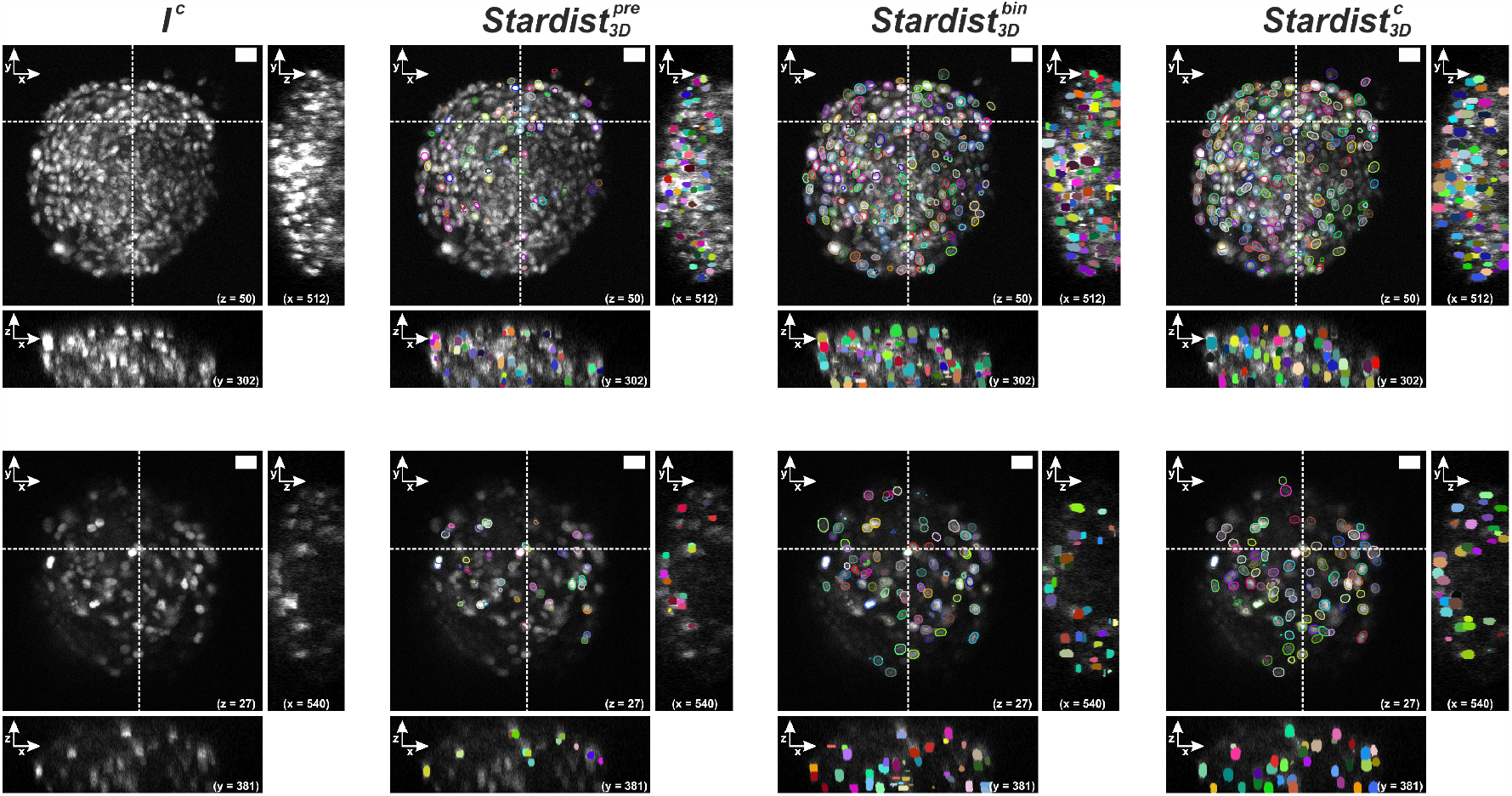
Comparison of segmentation obtained with 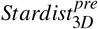 (Beghin et al., 2022), 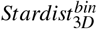and our 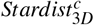 model trained with synthetic volumes.

**Figure 7:**
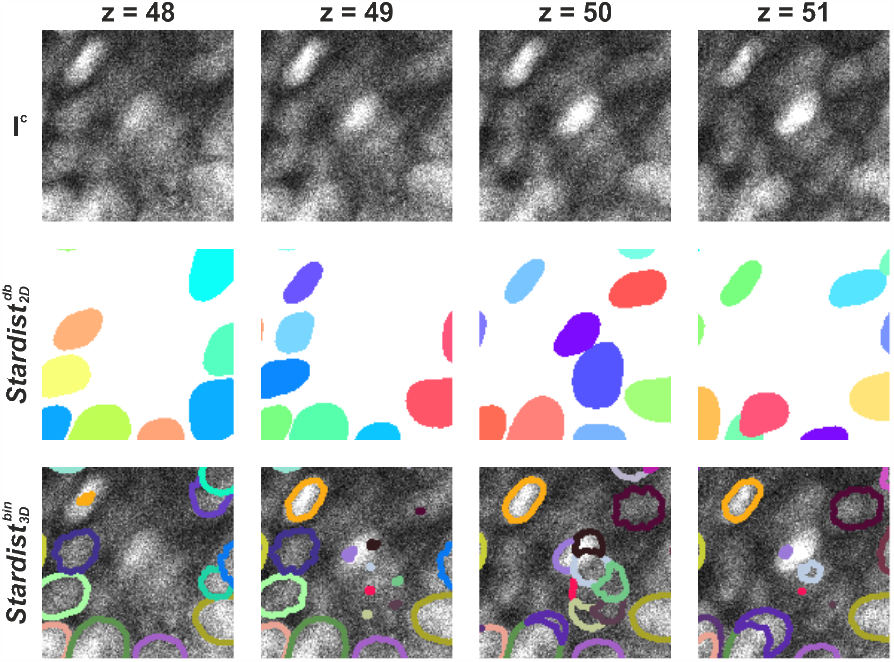
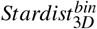 fails to properly handle isolated 2D masks by over-segmenting them.

### 4.2 Quantitative Assessment with *I*^*SD*^

Contrary to 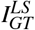 and *I*^*c*^ that exhibited cells having a bright and homogeneous texture, *I*^*SD*^ is composed of cells having a bright region surrounded by a dimer region which corresponds to the cell cytoplasm (Fig. 8left). Consequently, even if 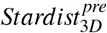 managed to identify more than 90% of the nuclei (Table 1), the overall segmentation is imperfect. While the model identified almost all the brightest nuclei (Fig. 8), explaining the high number of True Positives (TP), False Positives (FP) results from the model separating the bright and dim regions of a cell as 2 objects. False Negatives (FN), on the other hand, mostly originates from some nuclei exhibiting a different texture, for which the model only identified a small part of the membrane. 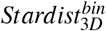 identified almost 99% of the nuclei present in *I*^*SD*^ (Table 1), thanks to 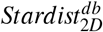 performing well in segmenting the nuclei in this low density nuclei distribution. Nevertheless, the segmentation being done in a per-frame basis, it led to the wrong identification of some cytoplasm as nucleus, explaining the large number of FP. Finally, we used our workflow to train a new Stardist 3D model 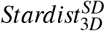 with paired of synthetic volumes 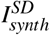 (generated by transferring the style of *I*^*SD*^ to the transformed masks 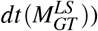) and masks 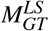. Similarly to 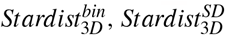 successfully detected 98% of the nuclei (Table 1). Nevertheless, directly accounting the axial direction for segmentation, in comparison to merging 2D masks to 3D as in 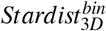, ended up with a lower number of errors with only 3 FP resulting from over-segmented cells.

**Table 1:** Comparison of the segmentation accuracy between 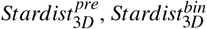 and 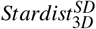.

| $I^{\text{SD}}$ | $\text{Stardist}_{3D}^{\text{pre}}$ | | | $\text{Stardist}_{3D}^{\text{bin}}$ | | | $\text{Stardist}_{3D}^{\text{SD}}$ | | |
| --- | --- | --- | --- | --- | --- | --- | --- | --- | --- |
|  | TP | FP | FN | TP | FP | FN | TP | FP | FN |
| Data 1 | 32 | 5 | 1 | 33 | 0 | 0 | 32 | 3 | 1 |
| Data 2 | 21 | 3 | 2 | 23 | 7 | 0 | 23 | 0 | 0 |
| Data 3 | 10 | 3 | 1 | 11 | 4 | 0 | 11 | 0 | 0 |
| Data 4 | 20 | 2 | 3 | 22 | 4 | 1 | 23 | 0 | 0 |
| Total | 83 | 13 | 7 | 89 | 15 | 1 | 89 | 3 | 1 |

**Figure 8:**
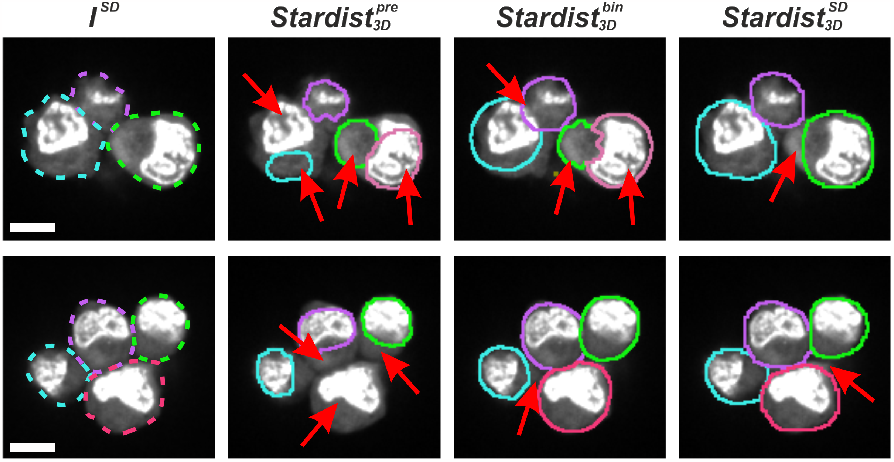
Comparison of the segmentation provided by 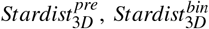 and 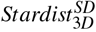, scale bar = 5 μm. Red arrows pinpoint regions with segmentation errors.

### 4.3 Discussion

Our workflow relies heavily on the FCGAN image synthesis capabilities which are, in our context, much more reliable than a CycleGAN thanks to the FCGAN’s multi-scale feature and paired training dataset. We have shown that using the pre-trained 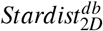 model directly on the acquired microscopy images and modifying the obtained instance masks was sufficient to properly train a FCGAN model. On the contrary, the segmentation provided by 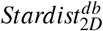 was not accurate enough to directly train a Stardist model in 3D such as 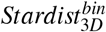. This discrepancy is related to the fact that 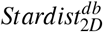 does not account for the axial direction, with several nuclei being therefore only segmented over one or two frames. It leads to a nuclei over-segmentation by 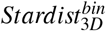, and therefore a possibly larger number of FP and FN. In our workflow, 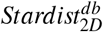 is only used to learn the style transfer. Creation of the synthetic volumes is done by applying the new style to 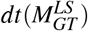, a dataset composed of hand annotated masks in which all the nucleus are accurately identified. Since our models 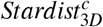 and 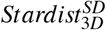 are trained with these synthetic volumes, they are mostly insensitive to the segmentation accuracy of 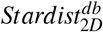 and are tailored to segment nuclei having the same texture as the acquisitions they have been trained on, therefore outperforming both 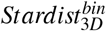 and 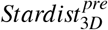.

Importantly, our synthetic volumes 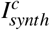 and 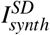 are composed of stacked generated 2D frames that does not guarantee intensity coherency in the axial direction (Fig. 9). They also lack smooth transitions between some appearing and disappearing nuclei as one would expect from a real acquisition. Nevertheless, our results confirm precedent findings (Baniukiewicz et al., 2019; Wu et al., 2023) for which training segmentation networks with synthetic volumes allows to achieve good performance.

**Figure 9:**
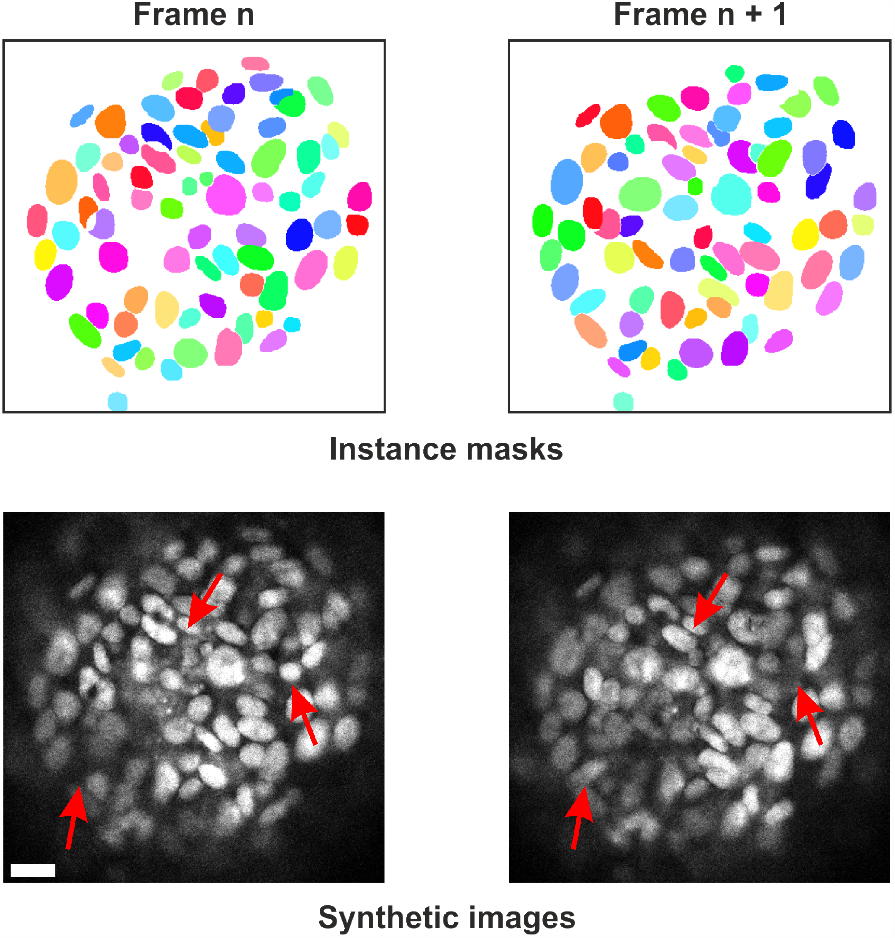
As synthetic volumes are composed of stack generated frames, a lack of smooth transition between appearing and disappearing nuclei is visible (red arrows), scale bar = 10 μm.

As already explained by the authors of Stardist (Schmidt et al., 2018; Weigert et al., 2020) and Cellpose (Stringer et al., 2021), we want to reinforce the fact that it is critical to train these models with data that have objects of similar sizes and features than the images we want to segment. Training Stardist with synthetic volumes as proposed in our workflow facilitates this process. It becomes indeed sufficient to resize 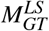 to generate images with nuclei exhibiting the same sizes and textures than the ones composing the acquired volumes we want to segment.

Finally, the border detection of the cells composing *I*^*SD*^ by 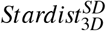 was not perfect and can be explained by two reasons (Fig. 8). First, the 3D segmentation provided by Stardist is highly convex while *I*^*SD*^ exhibits cells with irregular and possibly concave borders. Second, *I*^*SD*^ is composed of volumes with saturated intensity, resulting in a compressed dynamics. This combined with the small number of volumes prevented FCGAN to learn a perfect style mapping.

## 5 CONCLUSION

In this work, our objective was not to develop a method achieving state of the art accuracy for segmenting nuclei in 3D. In the past years (Mougeot et al., 2022), nuclei segmentation has seen the development of hundreds of methods competing for the leading position in term of segmentation accuracy. Unfortunately, most of these techniques are out of reach for life science labs and imaging facilities because they can be limited to one operating system or do not provide source code, tutorial or toy datasets (Mougeot et al., 2022). We therefore wanted to show that it is also possible to achieve good qualitative segmentation by using already established methods such as Stardist (Schmidt et al., 2018; Weigert et al., 2020) that are widely used in labs and facilities. Image synthesis with GANs is still, however, far from being easily accessible to life scientists. We therefore focused on a GAN architecture designed for the multi-scale organization of biological images. The FCGAN of Han (Han et al., 2020) was therefore an ideal choice as it allowed us to directly use our acquired microscopy images without cropping or finetuning parameters of the model.

All things considered, our workflow resulted in qualitatively good segmentation of microscopy volumes for which no GT existed. We were able to generate synthetic volumes having the style of different cell types and microscopy modalities from the same set of 3D GT masks (Fig. 10). Pairing them together, we trained several Stardist models tailored for each acquisition, managing to segment datasets without spending weeks in annotating volumes. We also quantitatively demonstrated that our workflow performed better than a pre-trained Stardist 3D model on a limited set of 4 volumes. In the future, we plan to push further this quantification as wells as further test its generalization capability by acquiring new datasets having a higher complexity than the *I*^*SD*^ dataset. We also plan to test Omnipose (Cutler et al., 2022) in our workflow, with the aim to better identify irregular cell borders.

**Figure 10:**
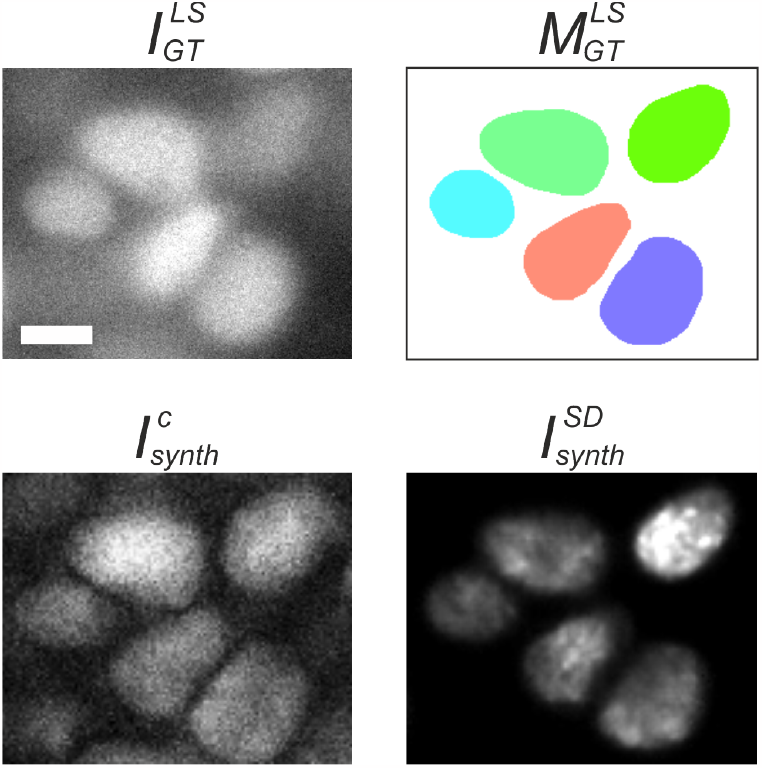
Generation of synthetic images from masks. From the same mask image (up), two synthetic images were generated with a different style (bottom).

Another point to consider is that the image synthesis provided by FCGAN will always be heavily dependent on the quality of the nuclei 2D segmentation. In our case, the pre-trained 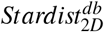 model gave satisfactory segmentation for feeding the FCGAN model. Otherwise, it will be necessary to generate new 2D manual annotations, a task still a lot easier than 3D annotations that can be accelerated by techniques such as SAM (Kirillov et al., 2023).

Finally, we think that this workflow could be improved by further manipulating the existing GT masks. GANs require similar distributions of the objects morphology and spatial organization between the source and target styles to generate realistic images (Liu et al., 2020). Similarly, Stardist would also benefit from training with pairs having the same distributions that the volumes to segment. This requires the ability to quantify the differences between these distributions, as it would allow to modify the GT masks to make them similar to the objects one would want to segment.

